# Unravelling the distinct phenotype and mechanosensitive properties of different tendon cell populations

**DOI:** 10.64898/2026.01.08.698354

**Authors:** S.E. Grossemy, D.E. Zamboulis, N.S. Khatib, M.R. Fazal, C.C. Gains, A. Giannopoulos, T. Hopkins, C Bevan, Y. Aggarwal, M.M. Knight, H.R.C. Screen

## Abstract

Tendinopathy arises from maladaptive cellular responses, though the drivers remain unclear. Here we identify and characterise a previously undescribed tendon cell population residing within interfascicular matrix (IFM), demonstrating its importance as the primary mechanosensitive cell in tendon.

We describe the first successful isolation and long-term culture of primary IFM and fascicular matrix (FM) cells, enabling direct comparison of their phenotypes and mechanosensitivity.

IFM cells exhibited a potent response to stiff substrates, displaying cytoskeletal remodelling, rapid drifting of tenogenic and ECM gene expression, and proliferative decline, while FM cells remained largely unaltered. Crucially, transferring IFM cells to compliant, IFM-like substrates recovered their proliferative capacity, morphology, gene expression.

This work defines IFM cells as the primary mechanosensitive tendon cell population, with implications for tendon ageing, injury, and regeneration. Importantly, it also enables identification of cell surface markers to isolate this population from other tendons, opening new avenues to explore mechanobiology-guided tendon therapeutics.

## Introduction

Tendinopathy encompasses a spectrum of prevalent tendon conditions characterised by chronic pain and functional impairment (1–3). Despite rising prevalence, effective therapeutic strategies remain elusive, largely due to incomplete understanding of the cellular mechanisms underlying disease progression. This knowledge gap largely reflects limited understanding of tendon cell phenotypes and their response to pathological changes in their microenvironment, such as tissue stiffening. A deeper understanding of tendon cell heterogeneity will enable development of physiologically relevant *in vitro* models, critical for elucidating disease pathways and developing targeted, effective therapeutic strategies.

Tendon is composed of highly ordered collagen fascicles, bound by soft, hydrated, predominantly non-collagenous interfascicular matrix (IFM). Several studies have identified the importance of IFM in tendon function, facilitating fascicle shearing to distribute strain through tendon (*5–7*). Critically, recent work has identified that distinct cell populations reside within fascicular matrix (FM) and IFM. Single-cell transcriptomics of murine, equine and human tendon cells indicate clustering of distinct cell subtypes, suggesting independent roles for each population in physiology and disease (*13–15*). However, relating these data to the tendon niche environments and elucidating their functional roles remains challenging.

The IFM niche is highly cellular, and *in vivo* and explant studies suggest it contains a more metabolically active cell phenotype (*8–10*) relative to elongated and fibroblastic FM cells (*11*). There is evidence that IFM cells specifically may drive tendon inflammation and degeneration (*10*). However, this is inferred from *in situ* studies, as there are no established protocols to separate IFM and FM cells, to interrogate their phenotype or response to pathology-driven environmental alterations (*12*). Without defined population-specific tendon cell markers, there is no solution to differentially isolating and studying these cells. It is crucial we elucidate both the phenotype and *in vitro* maintenances approaches for IFM and FM cells if we wish to build predictive *in vitro* disease models from which mechanistic understanding of tendon disease and therapeutic testing can be enabled.

With FM cells aligned along relatively stiff collagen fascicles (figure 1a), and rounded IFM cells residing in the soft gel-like IFM, it is hypothesised that these distinct cell types require niche-specific physical microenvironments to retain their characteristics. Injury and ageing both stiffen IFM (*5*), however it is currently unknown how this impacts tendon cells. Stiffening has been hypothesised to augment tendinopathy risk, potentially impacting IFM cell function and phenotype through aberrant mechanobiological pathway activation (*16*). Stiffening could initiate a feed-forward loop in which stiffer IFM alters ECM secretion, leading to further stiffening, progressing disease (*16*). Understanding how distinct tendon cell subpopulations respond to stiffness changes offers clear potential for unravelling the mechanisms behind ageing and tendinopathy.

**Figure 1:**
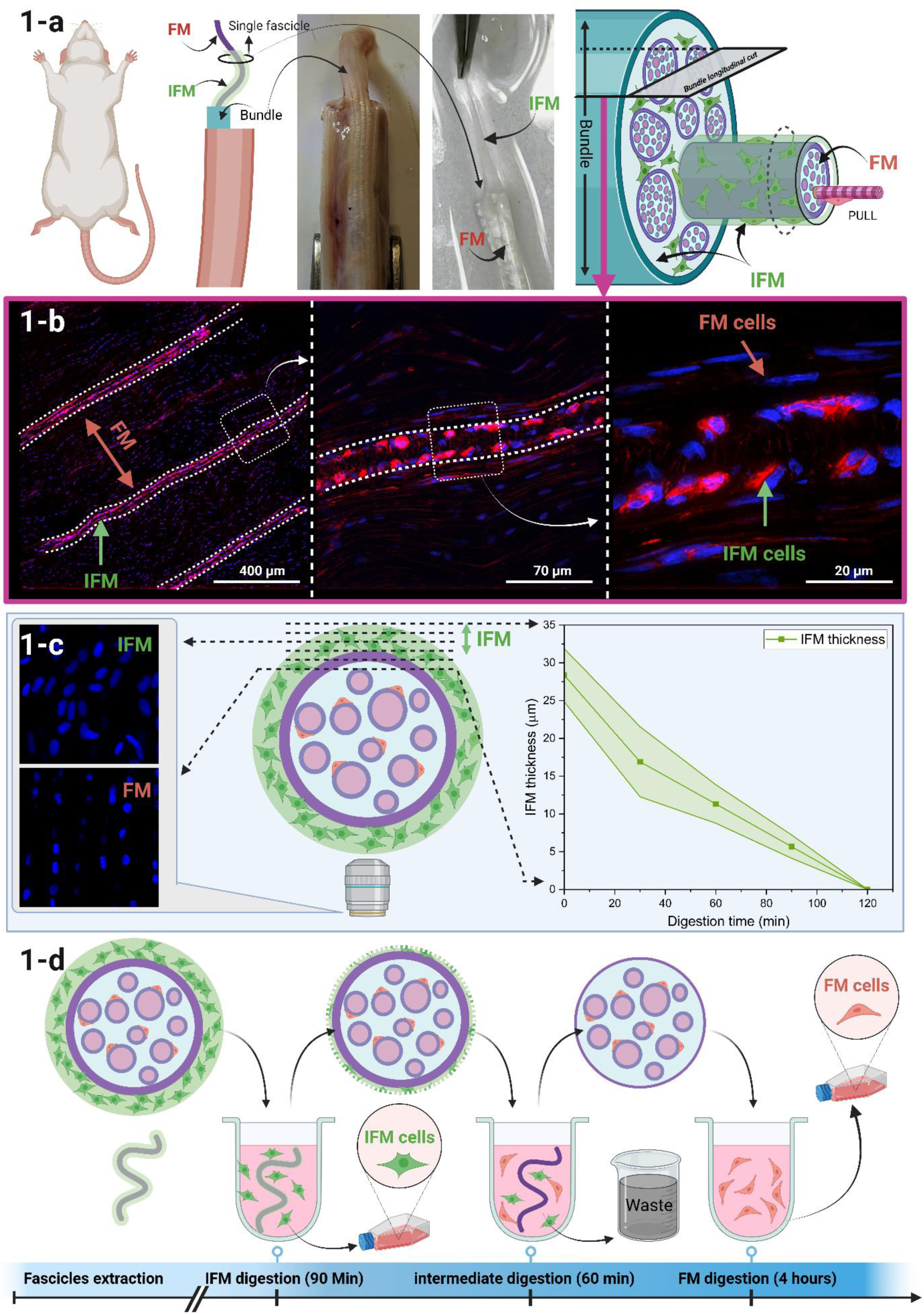
Rat tail fascicle sequential digestion allows separation of tendon sub cell populations. a) Schematic and photographic overview of the rat tail anatomy highlighting the location of interfascicular matrix (IFM) and fascicular matrix (FM) regions within the tendon, followed by a 3D schematic of bundle structure. b) Immunofluorescence of a rat tail quadrant (longitudinal cuts) stained with DAPI (nucleus) and phalloidin (actin cytoskeleton). IFM and FM regions are visually evident with different morphology and organisation of IFM and FM cells. c) The schematic depicts a transverse plane through an isolated fascicle surrounded by IFM as on removal from a tail tendon. Confocal images through the fascicle at different heights (shown on the left) demonstrate that it is feasible to detect when imaging within the IFM or FM region. Fascicles were digested in collagenase for 0 – 120 minutes, fixed, stained and imaged, and the thickness of the remaining IFM recorded at different time intervals, confirmed that IFM was consistently fully digested after 120 minutes. d) The digestion protocol was selected based on these data, discarding the digested cells collected between 90 minutes and 150 minutes to ensure IFM and FM digestions remained distinct for each cell population (n=3 rats, 2 fascicles/rat).

Here, we describe the first successful isolation of tendon IFM and FM cells, and novel techniques to maintain these distinct cellular phenotypes *in vitro*. Furthermore, we utilise these procedures to provide the first ever detailed phenotypic analysis of each cell population and identify new information about their relative mechanosensitivity. These transformative new techniques and analyses provide the crucial knowledge base to facilitate a step change in elucidating the drivers of tendinopathy.

## Results

### Sequential digestion successfully isolates tendon IFM and FM cells

We first established a ‘spatial separation’ approach for isolating IFM and FM cells. Rat tail tendon was deliberately selected due to its unique property in which individual fascicles with surrounding IFM sheath can be easily extracted without damage (figure 1a). A sequential digestion using collagenase and dispase was optimised, first digesting and extracting cells from the IFM, followed by further digestion and cell extraction from the FM (figure 1b-d).

An analysis of 30 fascicles (6/time point; 3 rats) demonstrated consistency across fascicles. Initial IFM thickness was 28.3 ± 3.6 μm and reduced linearly with time, with all samples showing some remaining IFM at 90 minutes, and none at 120 minutes. Based on these data, an isolation protocol was selected, in which IFM cells were collected from the first 90 minutes of digestion, FM cells from 150 minutes on, and digested material from 90-150 minutes discarded (figure 1d).

To validate IFM and FM cell separation, immunohistological observations of tenogenic and matrix markers *in situ* were cross-referenced with gene expression (RT-qPCR) in the isolated cells left to culture for 6 days (defined as Passage 0; P0) (figure 2a & 2b). Markers for each population were selected based on previous single cell sequencing work in which IFM and FM cell cluster gene expression profiles were reported (*14*).

**Figure 2:**
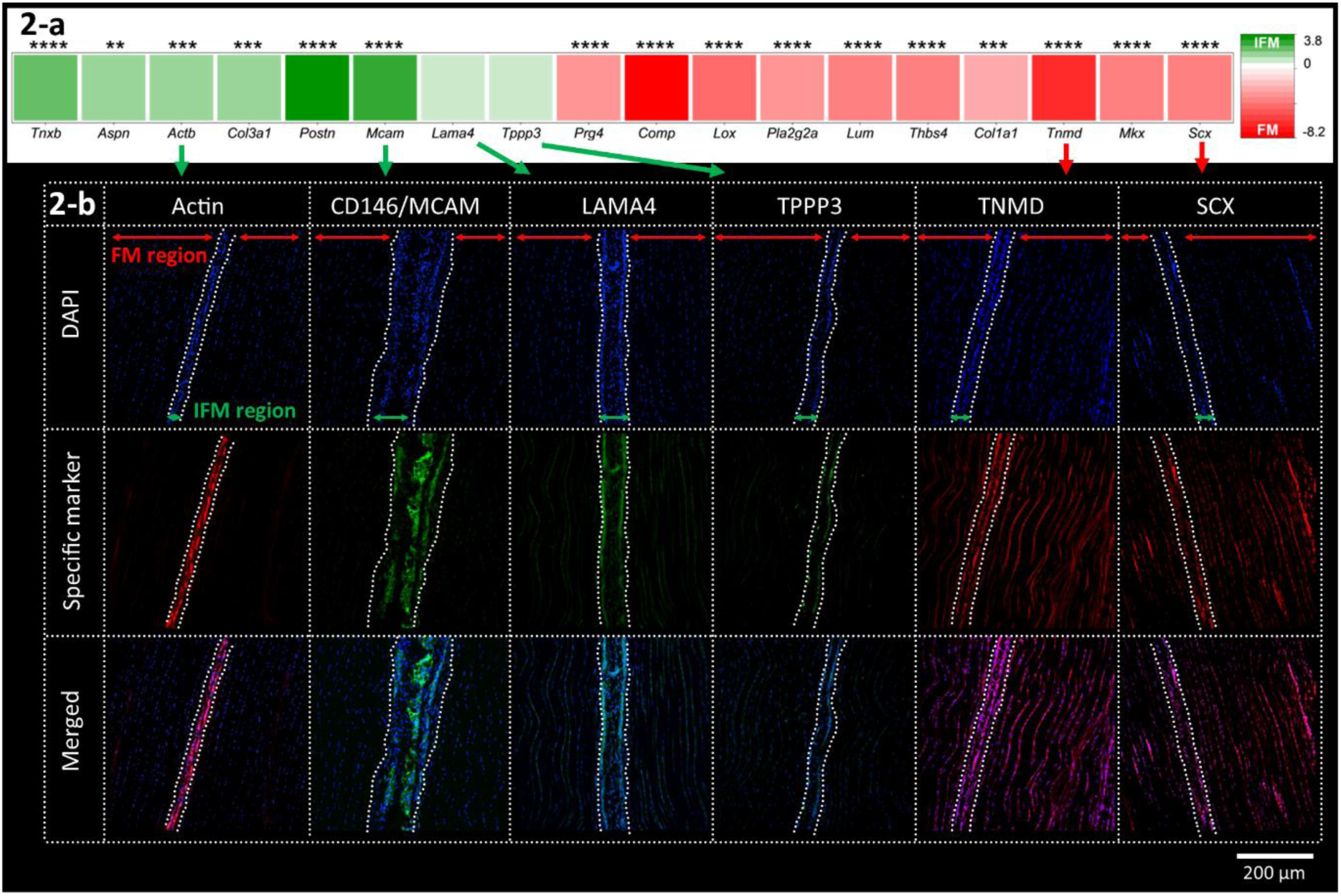
IFM and FM cells separated using sequential digestion exhibit distinct gene expression patterns reflective of those seen in their in situ niche. a) Heatmap showing RT-qPCR expression of key IFM and FM cell genes in isolated populations. Values represent the log₂ fold change of gene expression in IFM relative to FM cells, normalised to Gapdh. A value of 0 indicates equal expression, green indicates enrichment in IFM cells, and red indicates enrichment in FM cells. (*= p<0.05, **= p< 0.01, ***=p<0.001, ****= p<0.0001) b) Immunofluorescence of transverse rat tail tendon sections in which the IFM region is denoted between the white dotted lines. Cell nuclei are shown with DAPI staining (top row), followed by staining for specific markers (middle row). Actin, Cd146, lama4 & Tppp3 are localised to the IFM, whilst tenomodulin & scleraxis are localised to the FM region.

Gene expression profiles demonstrated clear markers for each cell type, corroborated by colocalised spatial protein expression in each compartment. ECM coding genes tenascin X (*Tnxb*), asporin (*Aspn*) and type III collagen (*Col3a1*) were all significantly more expressed in IFM than FM cells (p<0.01-0.0001). IFM cells also more highly expressed actin (*Actb*), Melanoma Cell Adhesion Molecule (*Mcam*; also known as *Cd146*), and periostin (*Postn*) (p<0.001-0.0001). Immunofluorescence confirmed localisation of actin and MCAM/CD146 to the IFM region. Laminin (LAMA4) and Tubulin Polymerization Promoting Protein Family Member 3 (TPPP3) tended towards greater gene expression in IFM cells and demonstrated spatial localisation to the IFM region.

In contrast, tenogenic markers tenomodulin (*Tnmd*), scleraxis (*Scx*) and mohawk (*Mkx*) exhibited significantly higher expression levels in FM cells (p<0.0001), whilst spatially localising to both the FM and IFM regions. ECM coding markers type I collagen (*Col1a1*), lubricin (*Prg4*), lumican (*Lum*) and cartilage oligomeric matrix protein (*Comp*) were more highly expressed in FM cells (p<0.001-0.0001). Finally, secretory phospholipase A2 group IIA (*Pla2g2a*) and thrombospondin-4 (*Thbs4*), which were specific to FM clusters in other studies, were more highly expressed in FM cells (p<0.0001).

Our findings demonstrate a robust sequential digestion approach for isolating FM and IFM tenocytes and retaining their phenotype for 6 days of culture. Heterogeneity between IFM and FM cell populations was confirmed, and notably, 15 of 18 genes showed consistent IFM/FM cell expression patterns to those seen in equine SDFT cells assessed with scRNA sequencing (14)(figure S8).

### IFM and FM cells show distinct responses to cell culture on tissue culture plastic

We next cultured IFM and FM cells on tissue culture plastic (TCP) for 4 passages (P0 – P3), exploring cell proliferation, morphology and gene expression across passages and time points (n=4; figure S1).

There was an observable increase in IFM and FM cell numbers during P1. However, during P2 and P3, IFM cells only maintained numbers while FM cells continued to proliferate (figures 3a and S2). Quantification of DNA content (Hoechst assay) at each time-point confirmed significant growth of both IFM and FM cells across 4 days at P1 (p<0.001), but by P3, whilst FM cell numbers increased >800% across 4 days (p<0.001), IFM cell number showed no increase over this time frame (figure 3b). These findings were corroborated with a haemocytometer approach to cell counting (figure S2).

**Figure 3:**
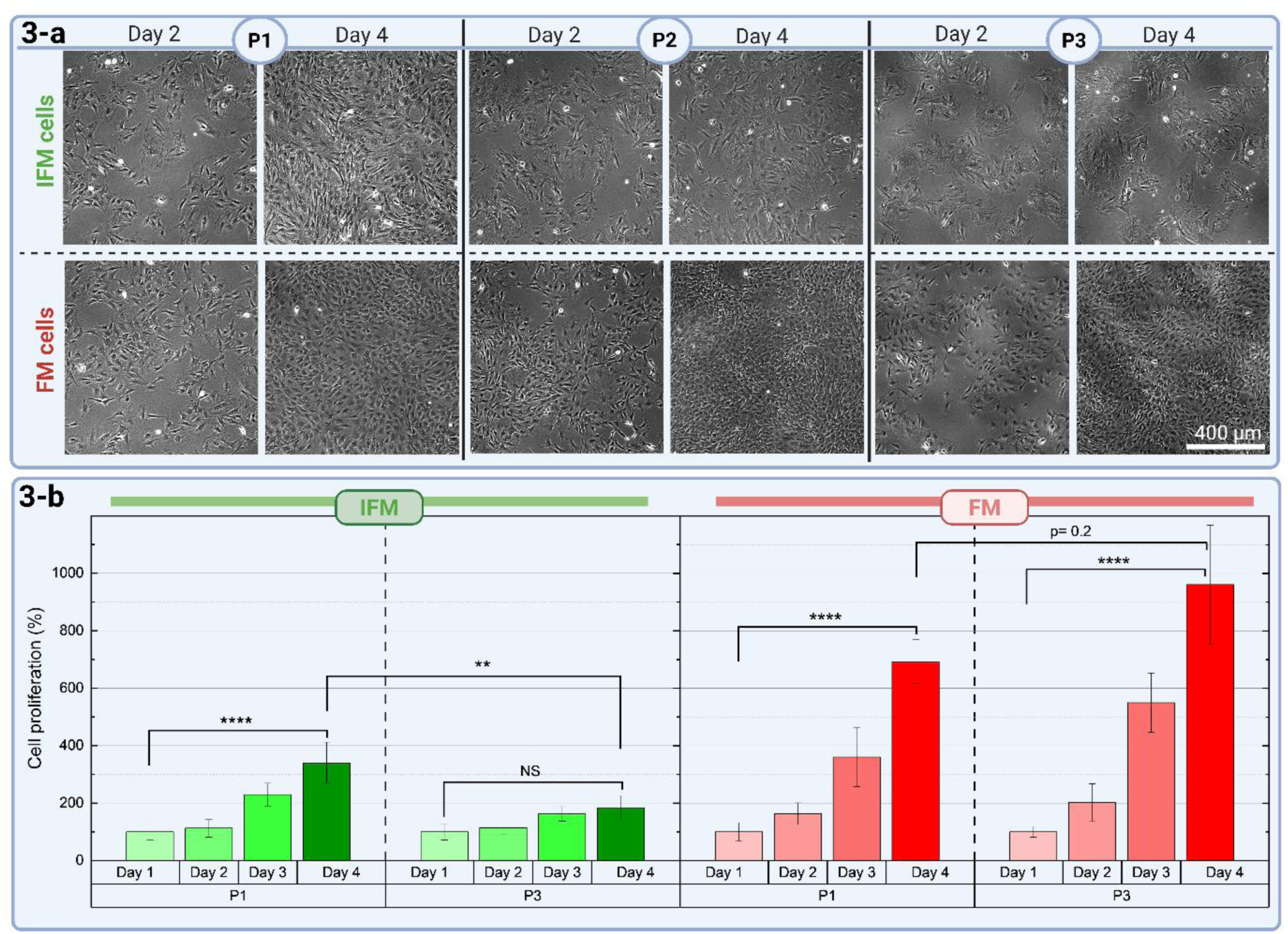
Serial passage on rigid tissue culture plastic selectively suppresses IFM proliferation but not FM cell growth. a) Phase contrast imaging of IFM and FM cells cultured on tissue culture plastic. b) Percentage increase in cell numbers as quantified with the Hoechst DNA assay over 4 days of growth following passage 1 and 3 (mean ± S.D.). **p<0.01, ****p<0.001.

Visual observation revealed that IFM cells substantially increased size and upregulated stress fibre formation across the cytosolic area from P0 to P3 (figure 4a). Quantification of cell body area, nucleus area, actin expression, cell eccentricity and alignment (>200 cells; n=4; Cell profiler (*17*)) focused on day 2 data, as FM cells were too confluent for analysis at day 4. IFM cells were significantly larger than FM cells at all passages (p<0.0001). In addition, whilst FM cell body area remained consistent across passaging, IFM cell area grew significantly with each passage (p<0.0001, figure 4b). Notably, nucleus area was significantly larger in FM than IFM cells at P0, but whilst nucleus size increased in both cell populations, this increase was far more pronounced in IFM cells, such that the IFM cell nucleus was significantly larger than that of FM cells by P2 (p<0.0001, figure 4b). Actin fluorescence/cell was significantly higher in IFM than FM cells at all time points. No notable differences in FM cell actin expression were evident with ongoing culture, whilst P3 showed significant (8-fold) increases in IFM cell actin compared to P0 (p<0.0001, figure 4b). Collectively, data highlights notably more pronounced changes in IFM cell morphology and actin fluorescence with TCP culture than evident in FM cells (figures 4b, S6).

**Figure 4:**
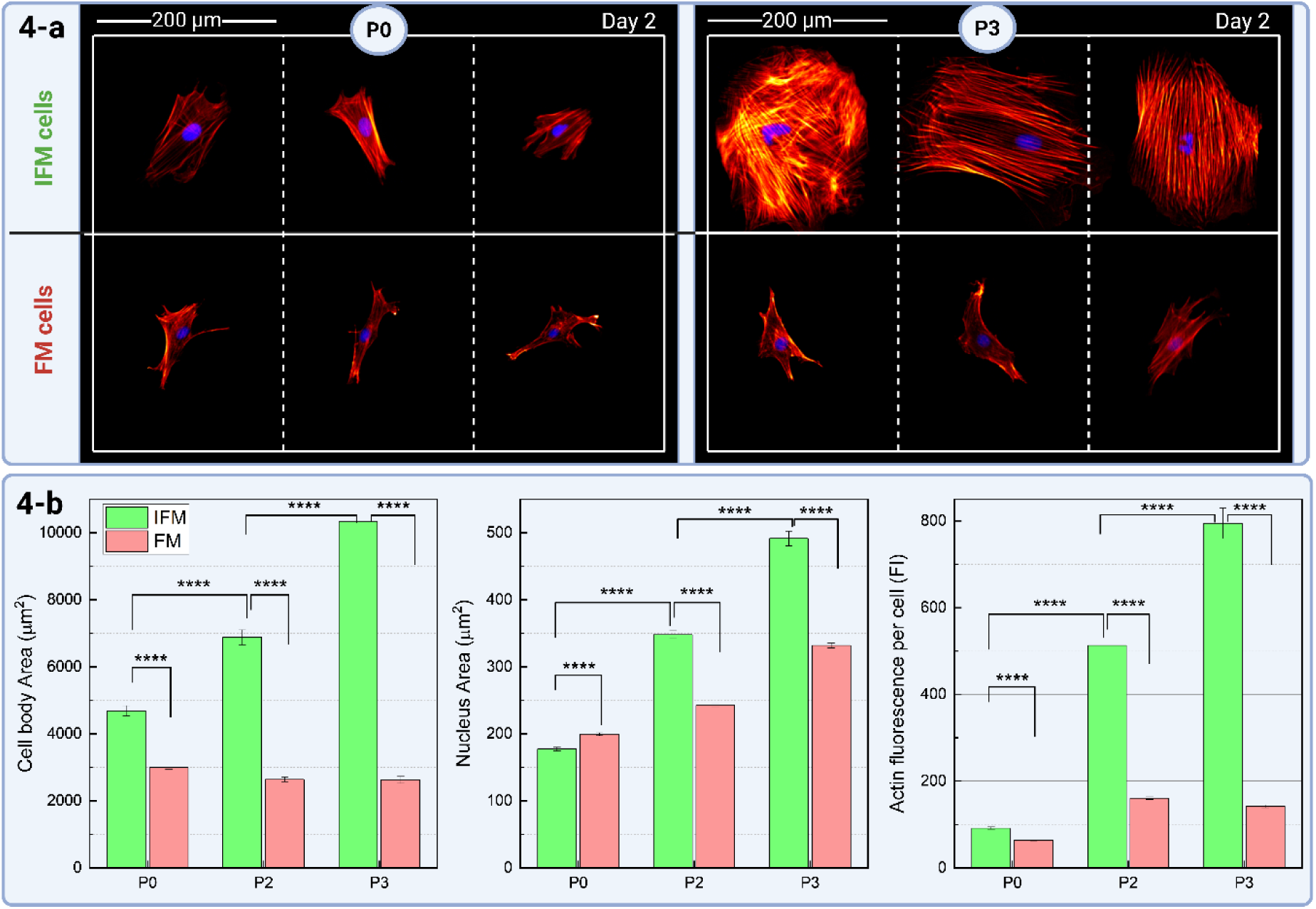
IFM cells exhibit enlargement and actin stress-fibre formation with passaging on tissue culture plastic, while FM cells remain consistent. a) IFM and FM cells fixed and stained with DAPI (nucleus) and phalloidin (actin), comparing cell morphology at day 2 of P1 and P3. b) Comparison of mean cell area, cell nucleus area and actin expression/cell at P0, P2 and P3 (N=200/group, mean ± S.E.), comparing IFM cells (green) and FM cells (red). ****p<0.001. Further associated data is provided in figure S6.

Finally, gene expression changes for IFM and FM cells were explored (RT-qPCR), comparing P2 and P3 with P0 (figure 5). Gene selection followed that previously quantified at P0 (figure 2a), based on the most commonly investigated tenogenic markers (*18, 19*) and a number of matrix proteins that localise to IFM or FM regions (*8, 20*).

**Figure 5:**
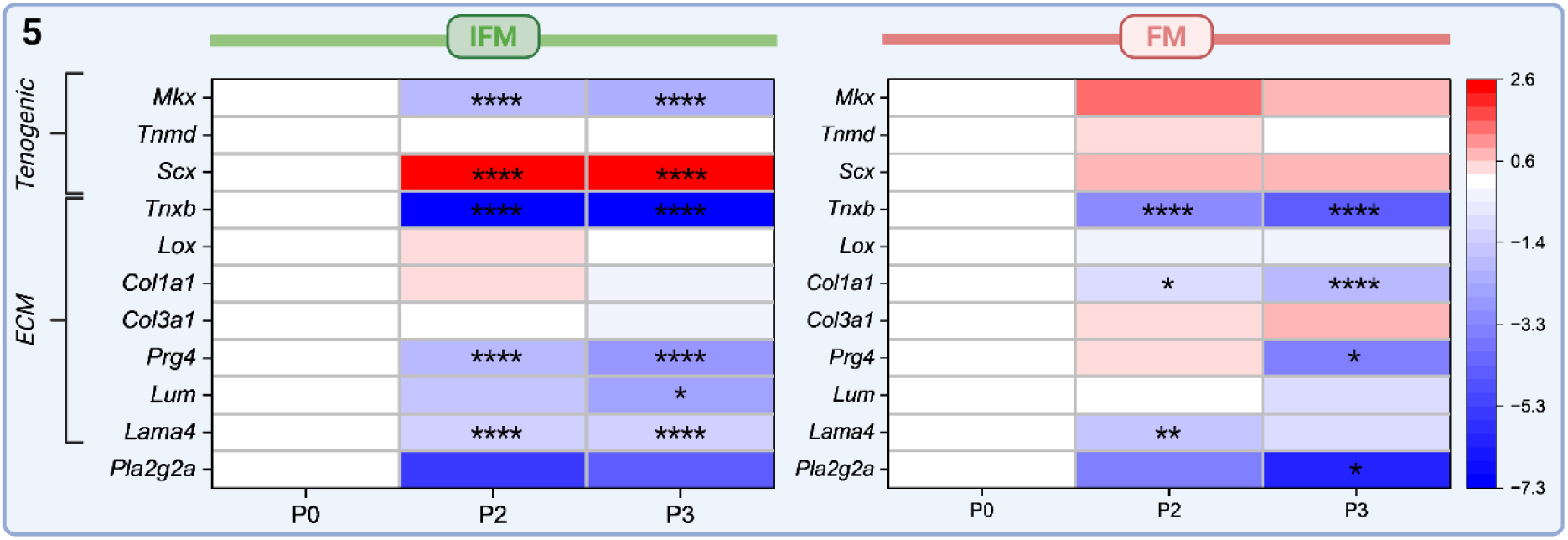
Culture on tissue culture plastic induces transcriptomic drift in both IFM and FM cells, but more potently in IFM cells. RT-qPCR of IFM and FM cells, exploring expression of a selection of genes through passages P0, P2 & P3. Expression values are presented as −ΔΔCT, which reflects the log₂ fold change of P2 and P3 relative to P0. The heatmap is centred on zero (white = no change from P0), with red indicating upregulation and blue indicating downregulation. Statistical comparisons are made relative to P0 for each gene and cell type. n=4/group. *p < 0.05, **p < 0.01, ***p < 0.001, ****p < 0.0001.

Genes encoding for tenogenic markers (*Scx*, *Tnmd, Mkx*) were all more highly expressed in FM than IFM cells at P0 (figure 2a). Following passages, *Mkx* expression significantly reduced in IFM cells (p<0.0001), but showed non-significant changes in FM cells (figure 5). *Tnmd* expression remained relatively stable across both cell types and passages, while *Scx* expression was upregulated in IFM and FM cells, but only significantly in IFM cells (p<0.0001).

For ECM related genes, both IFM and FM cells exhibited significantly reduced *Tnxb* expression by P2 (p<0.0001), with changes more pronounced in IFM cells. P0 data demonstrated that *Col1a1* was initially more highly expressed in FM than IFM cells, and vice versa for *Col3a1* (figure 2a). Following passaging, collagen expression remained broadly consistent in IFM cells, however FM cells transition away from *Col1a1* expression (p<0.0001), towards increased *Col3a1* expression (figure 5). Expression of glycoproteins (*Prg4, Lum, Lama4*) all significantly reduced with passage in both IFM and FM cells (p<0.05-0.0001, figure 5), except for *Lum* in FM cells. However, a more potent decrease in *Prg4* and *Lama4* expression was evident in IFM cells (p<0.0001). Collectively, while there was evident phenotypic drift in both cell types cultured on TCP, changes in IFM cells were more rapid and substantial in terms of both the range of genes and intensity of fold changes.

Together, data highlight more rapid phenotypic drifting of TCP cultured IFM-derived cells than FM cells. IFM cells lose their characteristics when cultured beyond P2 (or 12 days), a critical factor of consideration when developing *in vitro* tendon models (figures 3a,b, 4a,b & 5).

### IFM cells recover their original phenotype and activity when cultured on softer substrates

As IFM is significantly softer than FM, the pronounced IFM cell phenotypic drift on TCP may arise from the more extreme change in their physical environment. To explore phenotypic plasticity and mechanosensitivity further, we investigated whether IFM and FM cell phenotype could be recovered by transferring the cells to softer surfaces following drifting. Cells were grown on TCP until P2 and then at the following passage, either maintained on TCP (2.3-3.3 GPa stiffness), transferred to collagen coated TCP (2.3-3.3 GPa), or transferred to collagen-coated PDMS surfaces manufactured with various stiffnesses of 2.5 MPa, 900 kPa, 90 kPa or 20 kPa (figure S3a) to explore changes in cell proliferation, morphology and gene expression as previously described (n=4; figure S3b). Additional cells were transferred to these surfaces at P1 to explore proliferation rates. Each stiffness was quantified in figure S4.

Day 4 images of P3 cells demonstrated collagen coating did not improve IFM cell proliferation on TCP (figure 6a). However, IFM cells transferred to softer collagen-coated surfaces recovered their proliferation rates. DNA quantification confirmed IFM cell proliferation rates significantly increased with reducing substrate stiffness, and on the softest substrate (20 kPa) was significantly higher than proliferation on TCP (p<0.01, figure 6b). No significant changes in proliferation rates were evident for FM cells on any substrate (figure 6b). Comparing the impact of transfer to softer surfaces at P1 or P3 showed no significant differences in IFM cell proliferation rate between P1 and P3 transfer on any soft PDMS surface, but FM cells consistently proliferated faster at P3 than P1 transfer (figure S5). It is of note that despite significant proliferative decline, IFM cells transferred to soft surfaces at P3 still restored their proliferative function to that seen with P1 transfer (figures 6a,b, S5).

**Figure 6:**
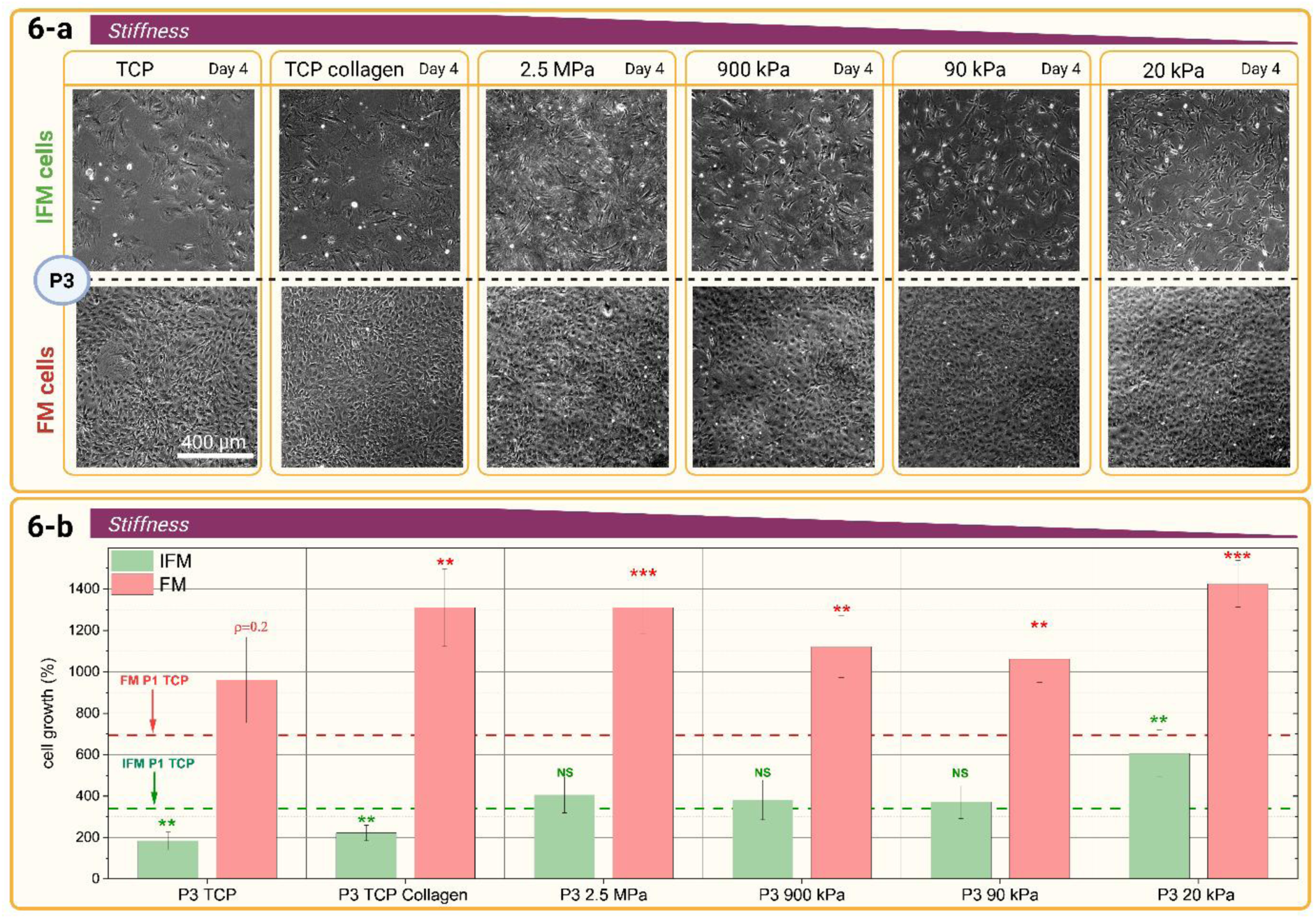
IFM cell proliferative capacity is restored in subsequent culture on soft substrates. a) Typical phase contrast microscope images of P3 IFM and FM cells on each substrate after 4 days of culture, highlighting the increased proliferation of IFM cells on softer substrates. b) DNA quantification for cells culture over 4 days at passage 3 (P3) across a range of substrate stiffnesses. Values are expressed as percentage growth relative to Day 0 (TCP) within each experimental replicate (n = 4). IFM cells show increased proliferation on softer substrates, particularly 20 kPa, whereas FM cell proliferation is high on TCP and largely unaffected by substrate stiffness. Dotted red and green lines indicate mean proliferation levels at Day 4 for P1 FM and P1 IFM cells on TCP, respectively, providing a reference for assessing phenotypic recovery or drift. Significance is calculated across conditions for each cell type. *p < 0.05, **p < 0.01, ***p < 0.001, ****p < 0.0001. full data set presented in S5.

Morphological observations further highlighted the dose-dependent effect of substrate stiffness on IFM cells only. P3 IFM and FM cells were fixed and stained at day 2 and 4, on each stiffness surface. FM cells had typically reached confluence by day 4, therefore semi-quantification focus on day 2 analysis, with data from day 4 only shown (figure S6) where individual cells could be assessed.

Representative IFM and FM cell images across substrate stiffnesses highlight no changes in FM cell size, but a potent reduction in IFM cell size as substrate stiffness reduced towards more physiologically representative values (figure 7a). Quantification of >200 cells demonstrated significant reduction in cell body area, nucleus area and total actin fluorescence in IFM cells with clear correlation between decrease in substrate stiffness and each parameter (p<0.001, figure 7b). Comparing P0 IFM cells to P3 IFM cells on the softest surface revealed that cell body size recovered by day 2 of culture, while nucleus size and total actin fluorescence only recovered by day 4 (p<0.001, figure S6).

**Figure 7:**
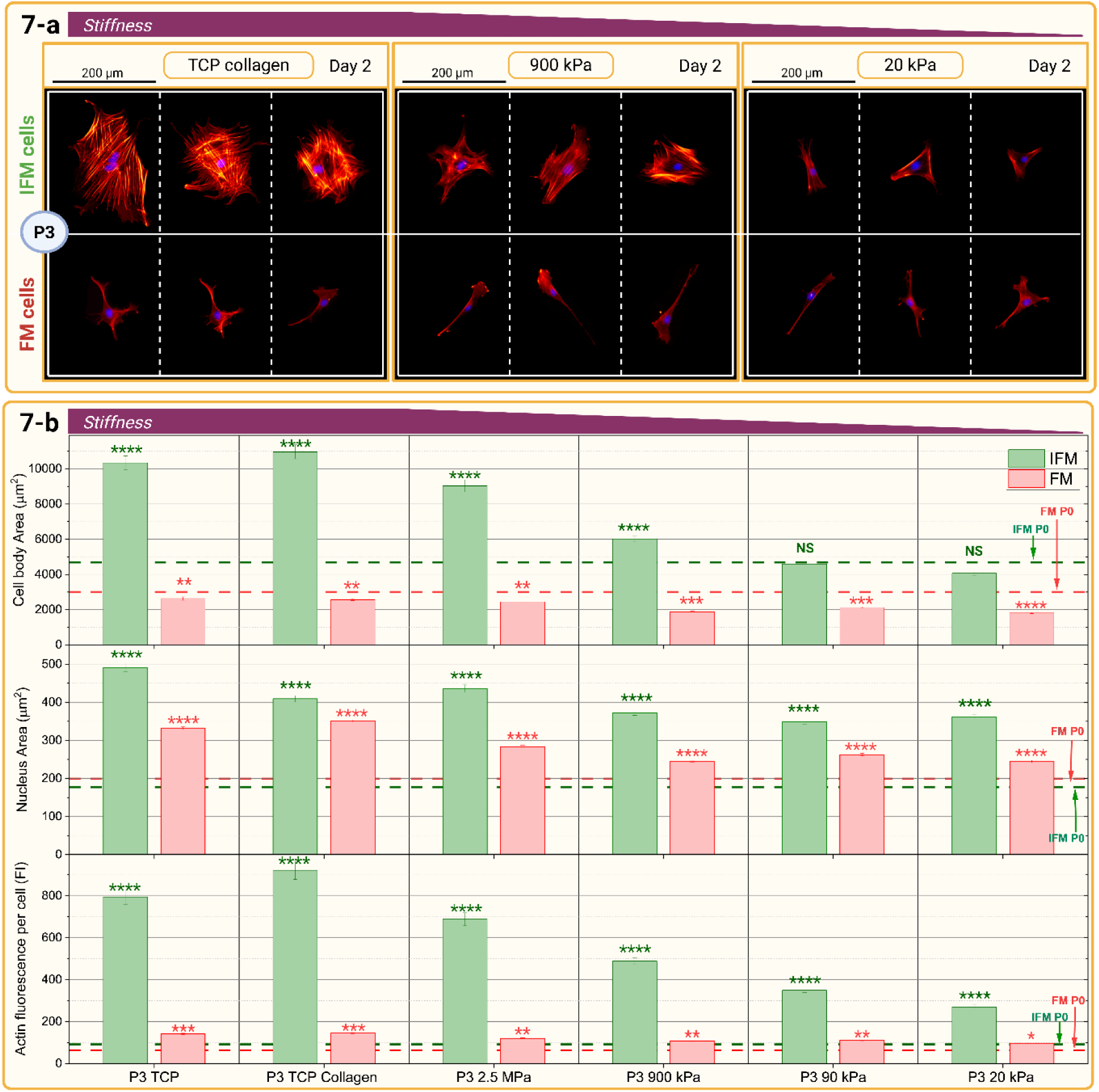
Culture on soft substrates recovers IFM cell morphology, while FM cell morphology is unaltered. a) Typical images of IFM and FM cells at day 2 of P3 culture comparing collagen-coated TCP with 900 kPa and 20kPa surfaces, illustrating the morphological recovery. Cells are stained with DAPI (nucleus; blue) and Phalloidin (cytoskeleton, red). b) Image semi-quantification of changes in cell body area, nucleus area, and actin expression/cell at P3 of culture on substrates of varying stiffness. Mean ± S.E. of >200 cells/condition are displayed for IFM cells (green) and FM cells (red). *= p<0.001 highlights a significant difference between the condition and the P0 TCP reference. A more detailed breakdown of the distribution of cell morphologies for each condition and correlations between cell body area, nucleus area and actin expression/cell are shown in figure S6&S7.

Finally, IFM and FM cell gene expression following P3 transfer to softer substrates explored whether the tenogenic phenotype can be recovered (figure 8). RT-qPCR at day 6 of culture on each surface explored the same selection of tenocyte markers and matrix genes as previously selected (n=4, figure 5).

**Figure 8:**
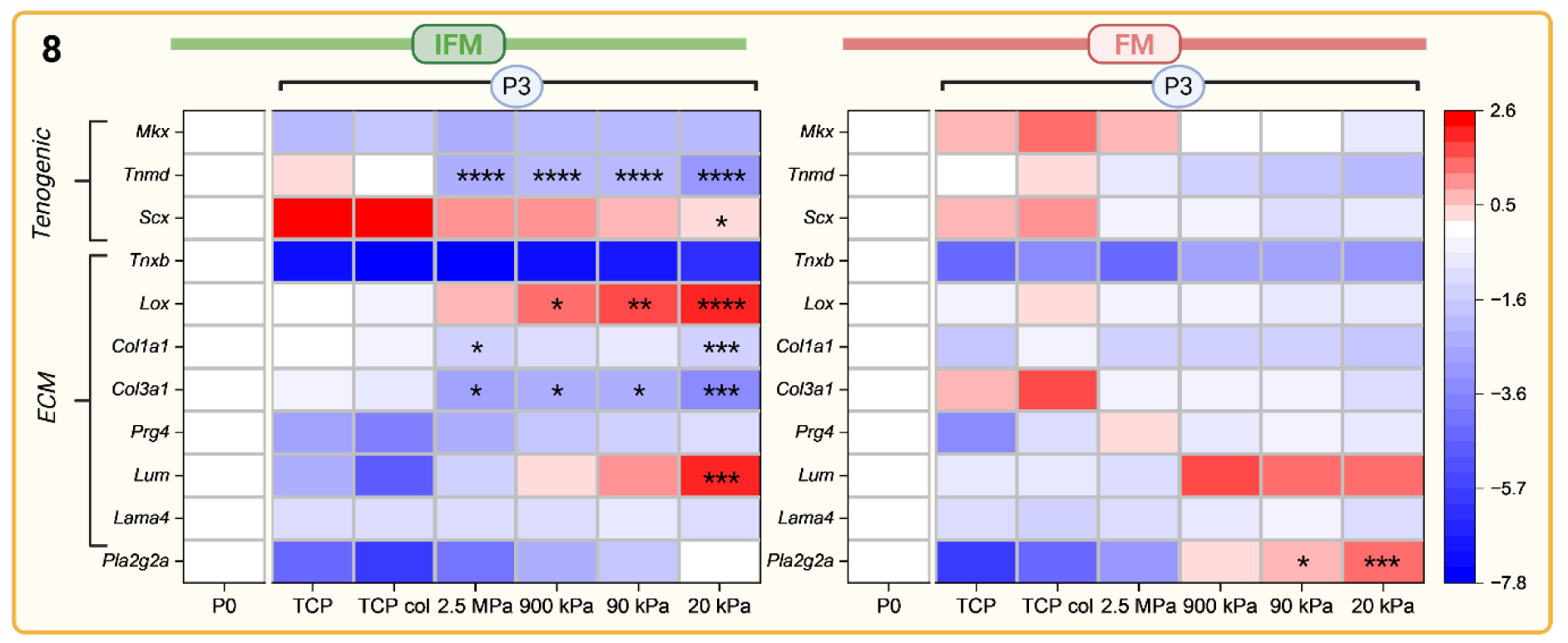
IFM cells exhibit pronounced gene expression shift in response to substrate stiffness, while FM cell gene expression remains relatively stable. In IFM cells, the majority of genes explored drift away from P0 expression during culture on TCP, and partially recover P0 expression when transferred onto softer surfaces. Expression values are presented as −ΔΔCT, which reflects log₂ fold change relative to P0. The heatmaps are centred on zero (white = no change from P0), with red indicating upregulation and blue indicating downregulation. Statistical comparisons are made relative to P3 TCP for each gene and cell type. *p < 0.05, **p < 0.01, ***p < 0.001, ****p < 0.0001.

Consistent with previous observations, the most significant gene expression changes with substrate stiffness were observed in IFM cells (figure 8). For tenogenic markers, culture on softer substrates led to dose-dependent downregulation of *Scx* towards P0 expression levels (p<0.05), and downregulation of *Tnmd* expression to levels below that observed at P0 (p<0.0001). In contrast, substrate stiffness did not appear to affect *Mkx.* For FM cells, only a tendency towards downregulation of *Mkx*, *Tnmd* and *Scx* was apparent.

Focussing on matrix gene expression, proteoglycans (*Prg4* and *Lum*) increased in both IFM and FM cells when cultured on successively softer surfaces, but changes were notably greater in IFM cells (IFM cell fold changes: 6-114; FM cell: 1.8-4.2; figure 8). In IFM cells, *Prg4* expression recovered towards P0 expression levels, while *Lum* showed significantly higher expression compared to P0 (p<0.001). In contrast, the expression of collagens (*Col3a1, Col1a1*) tended to reduce as substrate stiffness reduced, however only significantly for IFM cells (p<0.01-0.001, figure 8). Across all genes explored, the changes in gene expression with culture were consistently more pronounced in IFM cells, with most genes drifting away from P0 expression during culture on TCP and recovering back when transferred onto softer surfaces.

## Discussion

This study introduces the first definitive approach to isolate and maintain tendon cell subpopulations residing in the IFM and FM niche environments, and provides a first insight into their distinct phenotypes and properties. These outcomes make a step change in our ability to move beyond treating tendon cells as a single, homogeneous population, enabling crucial, more nuanced studies into the independent cellular drivers of tendinopathy. In addition, we have revealed the importance of the niche-specific mechanical environment in maintaining IFM and FM cell activity and phenotype. We provide evidence that IFM cells exhibit more pronounced mechanosensitivity to stiffness variations than FM cells, suggesting they are the primary mechano-responsive cell, with implications for their role in pathology and ageing.

Rat tail tendon cells were successfully separated into IFM and FM cells using a spatial sequential digestion to extract each cell population. FM cells expressed higher levels of tenogenic markers (*Scx*, *Tnmd, Mkx*), while also highly expressing *Thbs4* and *Col1a1*, in line with the higher abundance of COL1A1 in FM (Peffers et al., 2014). This finding is corroborated by a single cell RNA sequencing study of human hamstring tendon, which found that the MKX^+^ fibroblast cluster exclusively expressed *TNMD* and *THBS4*, and higher levels of *COL1A1* relative to other fibroblast clusters (Mimpen et al. 2024). Surprisingly, FM cells also expressed proteoglycan genes such as *Prg4* and *Lum* more highly that IFM cells, despite the IFM being more proteoglycan dense (Peffers et al., 2014). However, all human tendon fibroblast cell clusters have previously been shown to highly express genes related to proteoglycan production (Mimpen et al. 2024). As expected, IFM cells expressed higher levels of *Col3a1* than FM cells, a more abundant collagen in the softer IFM (Peffers et al., 2014). Interestingly, IFM cells also expressed higher levels of genes related to collagen fibrillogenesis (*Aspn*), fibril stability (*Tnxb*) and crosslinking (*Postn*), highlighting a potential role in regulating IFM collagen organisation, or possibly FM repair following fatigue or injury. This expression pattern was also found in an equine SDFT single cell sequencing study, whereby the cell cluster proposed as IFM fibroblasts expressed relatively higher levels of *Tnxb*, *Aspn* and *Postn* (*21*). Altogether, it is clear these two tendon fibroblast types show distinct phenotypes and roles, which should be considered in future *in vitro* experimental studies.

While it is notable that rat tail carries different functional demands to weight bearing tendons, we observed minimal cross-species differences in gene expression patterns between rat tail and equine SDFT cells. In rat tail, *Prg4* and *Lum* did not show the same higher IFM expression as seen in equine SDFT, but this may have arisen from the initial 6 days of TCP culture for rat IFM cells prior to analysis. The otherwise similar expression patterns across the two species is supportive evidence for conservation of IFM and FM phenotypes across species (figure S8). Importantly, this isolation approach provides a means to explore cell surface markers for distinguishing and isolating these subpopulations from other tendon digests. This finding, together with the clarity we provide on optimal cell maintenance conditions, opens opportunities to develop more comprehensive tendon *in vitro* models, considering cell heterogeneity when exploring drivers of disease and novel therapeutic targets.

Culturing IFM cells on TCP led to clear changes in cell proliferation, morphology and gene expression as early as P2. The effects of substrate stiffness on musculoskeletal cells are well-documented in the literature, and TCP known to strongly influence cellular behaviour and morphology (*22, 23*). Interestingly, FM cell behaviour remained relatively unchanged, apart from alterations in matrix gene expression. It is well established that intricate molecular control of cytoskeletal assembly, protein expression, and associated processes impacting cell phenotype in response to the mechanical environment, can differ significantly between cell populations (*24*), likely owed to varying levels of mechanosensor expression, such as integrins, focal adhesion kinase (FAK) and mechanically-sensitive ion channels. The difference in response to stiffness between the two cell types has implications for their relative roles in disease pathogenesis and progression. Further work exploring IFM and FM cell mechanotransduction pathway activation in response to stiffness may provide further clarity on the disparity of their response and effective mechano-therapeutic strategies to mitigate disease progression.

In a previous study investigating mesenchymal stem cell (MSC) culture, a cell type well known for its mechanosensitivity and phenotypic drift, responses to TCP culture were notably less pronounced than seen in IFM cells. Indeed, MSCs displayed a 1.5-fold expansion in cell body area from P6 to P9 and a 2-fold increase from P6 to P13 (*22*), relative to a 2.9-fold increase in IFM cell body area from P0 to P3, highlighting the remarkable responsiveness of IFM cells to substrate stiffness. The increase in IFM cell body area with progression through passages was directly proportional to actin expression, signifying a more extensive network of actin filaments, and stress fibres though passages. Notably, we observed what could be dorsal and ventral stress fibres and perinuclear actin caps developing. Perinuclear actin caps have been characterised as dense acto-myosin filaments specifically attached to the upper surface of the nucleus. They play a pivotal role in mechanosensation and mechanotransduction, enabling cells to sense changes in matrix compliance and respond to mechanical forces. Indeed, IFM cell body area and actin expression were clearly correlated to nucleus area. Stress fibres are wellknown to significantly impact nuclear shape, particularly through the actin cap, which compresses the nucleus and leads to an increase in its surface area (*25–27*). As passages progressed, stress fibres became more pronounced, causing the nucleus to flatten and the cell to spread across the substrate.

FM cells displayed less pronounced phenotypic alteration during TCP culture, but the increase in proliferation, minor morphological changes, reduced *Tnmd* and *Col1a1* expression and increased *Scx* expression though passages all suggest dedifferentiation. *Tnmd* and *Col1a1* are markers of mature tenocytes (*28*), whilst *Scx* is found in pre-differentiated tenocytes during development and has previously been determined as a specific marker of tendon progenitor cells (*29*). Further, other cell types, such as chondrocytes, Schwann cells and cardiomyocytes increase their proliferation rate when undergoing dedifferentiation (*30*). The increased *Scx* expression may also be directly linked to mechanical cues such as stiffness, as previous work demonstrated that *Scx* facilitates mechanosensing by regulating the expression of several mechanosensitive focal adhesion proteins (*31*). The intricate interplay between *Scx* expression, cell dedifferentiation, and mechanical cues demands further in-depth investigation.

It is of note that similar gene expression changes to those reported for FM cells have been observed when culturing tendon cells bulk-digested from human biceps tendons (*32*) and MSCs driven towards a tenogenic lineage (*33*). Viewed together with our data, this suggests that in conventional studies, where tenocytes are isolated by complete tendon tissue digestion, the digested cell population will undergo a rapid and substantial selection process, leading to dominance of FM cells.

To further explore cell plasticity, we transferred IFM and FM cells from TCP to softer substrates at P3. FM cells reversed the dedifferentiation changes seen with TCP passage, increasing *Tnmd* expression and reducing *Scx* expression in a linear manner with decreasing surface stiffness. Additionally, morphological observation revealed a tendency for FM cells to spontaneously align on surfaces with stiffness levels of 400 kPa and lower. Our data thus supports previous studies indicating that culture surfaces with a stiffness of 30-50 kPa are ideal for encouraging MSCs to differentiate down a tenogenic lineage or for tendon-derived cells to retain a tenogenic signature (*16, 34*).

Importantly, on the softest surfaces, IFM cells fully recovered original morphology and proliferation rate and showed a strong tendency to recover gene expression over six days. The 20 kPa stiffness was optimal for recovery, with a direct dose-dependent relationship between stiffness reduction and IFM cell recovery. It was notable that softer substrates promoted upregulation of genes associated with softer ECM components (*Lum, Prg4, Tnxb*), while stiffer substrates increased production of “stiffer” matrix components (*Col1a1, Col3a1*, *Lox)*. The intricate interplay between environmental stiffness and cell metabolism may contribute *in-vivo* to a detrimental cycle of tissue stiffening with ageing driving further matrix stiffening. Indeed, similar feedback loops have previously been reported in fibroblasts, where collagen production and *Lox* expression were upregulated on stiffer surfaces. Where those findings were translated *in vivo* they proved strongly involved in fibrosis for lungs, kidney, liver, and skin (*23*). Our previous research has reported that IFM stiffens with ageing (*35*), underscoring the relevance of our observations to broader physiological contexts.

Collectively, the heightened mechanosensitivity of IFM cells and their ability to adapt phenotype with environmental stiffness have significant implications for tendon research. The pathways and mechanisms modulated by substrate stiffness also mediate response to substrate stretch (*36*), building further evidence for a paradigm shift in our understanding of tendon disease, placing the IFM niche as the prime driver of tendon homeostatic maintenance and disease progression. Control of the IFM niche in tandem with cell therapies integrating the complexity and heterogeneity of the tendon cell populations, may provide avenues to rejuvenate IFM cells and provide advancements in tendon regeneration treatments.

## Materials and Methods

### Manufacture of PDMS substrates of different stiffnesses

To establish Polydimethylsiloxane (PDMS) substrates of specific controlled stiffness, Sylgard™ 184 and Sylgard™ 527 were used (Ellsworth Adhesives) following the manufacturers proprietary instructions. Sylgard™ 184 was prepared with a 1:10 ratio of curing agent to Sylgard™ 184. The components were rigorously mixed using a stripette for several minutes to ensure a homogenous solution. Subsequently, a vacuum was applied to remove entrapped air bubbles. For Sylgard™ 527, a 1:1 ratio of parts ‘A’ and ‘B’ was applied, with thorough mixing essential for uniform stiffness distribution.

To create substrates of varying stiffness, we combined Sylgard™ 184 and Sylgard™ 527 at the following ratios (184:527): 1:20, 3:20, 8:20 and 0:20. The mixture was vigorously stirred, then placed on a tube roller for 1 hour to homogenise.

### Compression test

PDMS samples for mechanical characterisation were prepared by casting into a metallic mould with a cylindrical hole measuring 5 mm in depth and 10 mm in diameter. A quasi-static Instron 3342, equipped with a 10 N load cell was used for testing. Unconfined compression was applied at a constant strain rate of 50%/minute, until a maximum compressive strain of 50% was reached. The compressive modulus of each sample was calculated from the gradient of the stress-strain curve between 0% and 10% compressive strain where the behaviour was linear.

### Preparation of PDMS surfaces for cell culture

Each PDMS mixture was dispensed into a 12 or 6 well plate, ensuring enough volume was added to fully coat the bottom of the respective well by gentle manual rotation of the plate. Plates were left undisturbed overnight at 60° C for curing, after which they were sterilised by exposure to 70% ethanol for 20 minutes, followed by three successive washes with sterile water for 5 minutes each.

Each PDMS surface was treated with a sulfo-SANPHA (Sigma-Aldrich, 803332) solution under UV light for 20 minutes at a concentration of 250 μg/mL in HEPES buffer (20 mM, pH 8.5) and then thoroughly washed three times with HEPES buffer. A solution of rat tail collagen type 1 (Merk, 08-115, diluted at 20 µg/mL in a 0.1% acetic acid solution) was then applied to the PDMS surfaces overnight at 4 °C, followed by three rinses with Phosphate-buffered saline (PBS) for 5 minutes each, after which treated surfaces were exposed to Dulbecco’s Modified Eagle Medium (DMEM) for at least 1 hour at 37 °C before cell seeding.

### IFM and FM cell extraction - protocol optimisation

Rat tails were from acquired from 11-week-old male Wistar rats (Charles River) for all experiments. To optimise the protocol, three tails were used. Each tail was skinned, sectioned into a proximal and middle section of ∼40 mm length each, discarding the remaining distal region. Fascicles were extracted from the proximal and middle sections by gently pulling individual fascicles from their sheath with forceps.

After extraction, fascicles were placed in a collagenase/dispase solution (2 mg/mL collagenase type II (Gibco, Thermo Fisher) and 1 mg/mL dispase (STEMCELL Technologies)) and incubated at 37° C with 5% CO_2_ with agitation (∼80 RPM). Two fascicles were removed from the digestion solution after 10, 30, 60, 90, and 120 minutes for each biological replicate (6 fascicles in total at each time point), washed in PBS, and fixed in 4% Paraformaldehyde (PFA) for 2 hours at room temperature. Fascicles underwent a further PBS wash, followed by two washes in PBS + 0.1% bovine serum albumin (BSA) before incubation in 4′,6-diamidino-2-phenylindole (DAPI) (1 μg/mL in PBS) for 10 minutes at room temperature in the dark. Following a final wash in PBS, the fascicles were mounted on glass for confocal imaging using a Leica Laser Scanning Confocal Microscope TCS SP2 at a x20 magnification (figure. 1c).

To establish the rate of IFM digestion, the thickness of the IFM layer at each digestion time point was measured. The surface of the fascicle was identified as the location where IFM cell nuclei were first seen as present on the top surface of the fascicle (*z*1), and from there, the confocal scanning depth progressively lowered into the IFM until the FM cell nuclei first became visible (*z*2). The distance between these two z-positions determined the thickness of the IFM layer surrounding the fascicles (*z*= *z*2-*z*1). Measurements were conducted on two regions along each fascicle length, and a mean value for IFM thickness was first determined for an individual fascicle, before a mean IFM thickness for each digestion time point was calculated across all 6 fascicles.

Based on our findings of IFM digestion rate, a sequential digestion protocol was optimised to ensure only IFM cells were present in the initial digest, and only FM cells in the final digest. Fascicles were digested for 90 minutes at 37° C in the collagenase/dispase solution with gentle agitation (∼80 RPM), to digest IFM tissue and release IFM cells (figure 1b). They were then removed from the digestion solution and rinsed in PBS for 5 minutes before transferring to a fresh collagenase/dispase solution for an additional 60 minutes incubation at 37° C with gentle agitation. On removal, fascicles were rinsed again for 5 minutes in PBS before submerging in a final fresh collagenase/dispase solution for 4-5 hours, until the remaining fascicular tissue was fully digested and the FM cells released.

The digest solution from the middle of the three digest periods contained a mix of IFM and FM cells and was discarded. The first and last digest solutions, containing only IFM or FM cells respectively, were passed through a 70 μm mesh filter to remove any remaining undigested matrix before the solution was centrifuged for 5 minutes at 300 g, the digest media discarded and the cell pellet resuspended in culture media (DMEM - low glucose, pyruvate, no glutamine, no phenol red, plus 10% FBS, 5% HEPES, 5% L-glutamine).

IFM and FM cells were cultured in T75 culture flasks at an initial seeding density of 7500 cells/cm^2^. Culture media was changed every 3 days, and after 6 days when confluency was reaching 90%, the cells were detached ready for further experimental use, using 2 gentle washes with 37° C PBS then a 5 minute exposure to a 0.01% trypsin 0.02% EDTA solution.

### Immunofluorescence of rat tail tendon

Full tendon bundles were isolated from the rat tails (n=2) to preserve the structural integrity of the interconnected fascicular matrix (FM) and interfascicular matrix (IFM) regions. The samples were embedded in OCT compound and snap-frozen in a bath of hexane surrounded by dry ice. Cryo-sections were prepared at a thickness of 15–20 µm and adhered to glass slides for immunofluorescence analysis. The cryosections were thawed before fixing with 4% PFA solution at room temperature for 10 minutes and then washed thrice with ice-cold PBS 4° C for 5 minutes. To stain for specific antibodies, sections were treated for 10 minutes with 0.1% Triton X-100 solution, washed three times in PBS, blocked with 5% goat serum and then incubated in the primary antibody overnight at 4° C. Following three rinses in PBS, secondary antibodies were applied at room temperature for 2 hours. The sections were washed thrice with PBS for 5 minutes before incubation in DAPI solution (1 μg/ mL in PBS) for 10 minutes and the three PBS washes repeated.

Stained sections were mounted with ProLong Gold Antifade Mountant (Thermo Fisher Scientific) and left to cure overnight. Images were taken using confocal microscopy (NIKON CSU-W1 SoRa) at 40x. Product codes and specific concentrations used for each antibody are specified in figure S4.

### Cell count though passages

Cell numbers were monitored at the end of each passage using a hemocytometer and trypan blue exclusion staining. At day 6 of each passage, cells were enzymatically detached using a trypsin-EDTA solution, centrifuged to form a pellet, and resuspended in culture medium. A 10 µL aliquot of the cell suspension was mixed with an equal volume (10 µL) of 0.4% trypan blue (Gibco, Thermo Fisher Scientific). The mixture was thoroughly homogenised by pipetting before loading 10 µL into the hemocytometer chamber. Only viable (unstained) cells were counted in four quadrants (top-left, top-right, bottom-left, bottom-right) using a light microscope, and the average cell count was used to determine the final cell number. Data was expressed as percentage cell growth, calculated as: Percentage Growth = (Final Viable Cell Count at Day 6 / Initial Seeded cell number Density) ×100. All cell counts and growth percentages presented in Figure S2 were normalised to the number of cells seeded/flask.

### Cell Proliferation using DNA quantification on different substrates

To assess cell proliferation rates on different stiffness substrates, IFM and FM cells from four biological repeats (n = 4) were used. Cells were seeded at a density of 2,500 cells/cm² for all conditions at P1 and P3, and proliferation was compared across collagen-coated PDMS surfaces at each stiffness and tissue culture plastic (TCP) with and without collagen coating.

A separate 12-well plate was prepared for each cell type (IFM and FM) and time point (day 1, 2, 3 & 4), such that 8 plates were prepared for each biological repeat, each containing a technical duplicate for each of the 6 stiffness conditions (TCP, TCP + Collagen, 2.5 MPa, 900 kPa, 90 kPa, and 20 kPa). This setup was replicated for each rat (n = 4).

At each collection time point, the culture medium was removed, and the samples were immediately stored at −80°C until further analysis. Phase-contrast imaging was additional performed at day 2 and day 4 to visually assess cell growth (DMI8).

For DNA quantification, frozen cells were lysed by adding 250 μL of Milli-Q water to each well, followed by a 1-hour incubation at 37°C and an additional 1-hour incubation at −80°C. A 100 μL aliquot of each solution was transferred to a 96-well plate in duplicate, and 100 μL of a 25 μg/mL working solution of Hoechst 33258 in TNE buffer (DNA Quantification Kit, Sigma) was added to each well. The plate was incubated in the dark at 37°C for 45 minutes before fluorescence readings were conducted using a fluorimeter (FluoStar) with excitation at 358 nm and emission at 461 nm. A standard curve was established for each set of readings using calf thymus DNA at five different concentrations. To account for variations in initial seeding numbers, cell counts for each population were normalised to the mean cell number on plastic at day 1 (100%).

### Cell Morphology quantification

Cell morphology was examined during drifting on TCP at day 2 and day 4 at P0, P2, and P3 in biological quadruplicate (n = 4), then across different stiffness substrates at P3.

In all instances, cells were seeded within 12-well plates with a single well allocated to each surface condition (just TCP or TCP + Collagen, or collagen-coated PDMS at 2.5 MPa, 900 kPa, 90 kPa, and 20 kPa), each cell type (IFM & FM) and each culture time (day 2 & day 4), for each biological repeat.

At each time point, the supernatant was removed, and cells rinse in DPBS (Ca+) at 37°C prior to fixing in a 4% PFA solution for 20 minutes. After three washes with ice-cold PBS for 5 minutes each, cells were exposed to Triton X-100 (0.1% in PBS) for 5 minutes and rinsed again with three PBS washes. Blocking was performed using a 1% BSA solution for 1 hour, before cells were stained with phalloidin (Texas Red™-X Phalloidin ThermoFisher diluted in 1% BSA solution) for an additional 45 minutes, followed by three PBS rinses. DAPI solution (1μg/ mL in PBS) was applied for 10 minutes, and samples were rinsed three times in PBS. The prepared samples were then ready for acquisition on an epifluorescence microscope (Leica DMi8) using a 20x lens. A minimum of 5 images/condition for each of the four biological replicates were taken. Morphological characteristics were quantified using CellProfiler, with a minimum of 200 individual cells analysed/condition. The extracted parameters included cell body area, nucleus area, integrated actin fluorescence intensity, cell eccentricity, and cell alignment.

DAPI staining was used to segment nuclei, while actin expression was used to define cell boundaries. Nuclei were identified as primary objects, and cell bodies were defined as secondary objects, ensuring each cell remained linked to a nucleus. Cells with segmented bodies touching image boundaries were excluded from the analysis to prevent artifacts. Image processing was performed using a custom CellProfiler pipeline, including segmentation, object classification, fluorescence intensity quantification, and morphological feature extraction. Final data were exported as CSV files for statistical analysis.

Data on morphological characteristics for all cells across all four biological repeats were combined to explore how morphology varied with passage and surface stiffness for both IFM and FM cells. Correlations between parameters for different cell passages or surface stiffnesses were also explored. To assess relationships between cell body area, nucleus area, and actin expression, a subset of cells was selected from each condition for analysis (1 image/rat). Scatter plots were generated to visualise the relationships between cell body area vs. nucleus area, actin expression vs. nucleus area, and actin expression vs. cell body area, separately for IFM and FM cells. For each pairwise comparison, the Pearson correlation coefficient (R) was calculated to assess correlation. Additionally, the M value was computed, representing the slope of the linear regression model fitted to each dataset.

### Gene expression quantification RT-qPCR

RT-qPCR analysis was conducted to assess the relative gene expression levels of tenogenic and ECM markers in IFM and FM cells. RNA samples were collected during drifting experiments at P0, P2, and P3, then at P3 during the recovery experiments across the range of stiffness surfaces (TCP, TCP + Collagen, 2.5 MPa, 900 kPa, 90 kPa, and 20 kPa). RNA samples at P0 were used to validate the cell isolation protocol, while other samples were used to evaluate the effect of long-term culture.

Three wells were seeded for each cell type (IFM and FM) in a 6-well plate, such that any test condition had a dedicated 6-well plate. Cells were seeded at 4,000 cells/cm², and after six days of culture, samples were collected for RNA extraction. The three wells for each cell type within a plate were pooled together to obtain sufficient RNA yield, resulting in one RNA sample/stiffness and cell type/rat.

Trizol (Invitrogen, 12044977) was applied directly to cell cultures, which were then stored at - 80 C° until subsequent analysis. Total cellular RNA extraction was performed using an RNA Miniprep Kit (Bioscience) according to the manufacturer’s instructions. The extracted RNA’s purity and quantification were determined by measuring absorbance with a NanoDrop 2000 spectrophotometer (ThermoFisher). To ensure uniformity, RNA samples were normalized in concentration before being transcribed into cDNA using Promega RT PCR reagents. For realtime PCR, 10 ng of RNA was utilized/reaction, with Sybergreen master Mix (Takyon). GAPDH was selected as a housekeeping gene to normalise all gene expression data.

### Statistical Analysis and Reproducibility

All data were obtained from four biological repeats (n = 4). Normality was assessed using the Shapiro-Wilk test for sample sizes below five and the Kolmogorov-Smirnov test for larger sample sizes. All data were found to be normally distributed, therefore parametric tests were applied throughout. Statistical analyses were conducted using the OriginPro software, with a significance threshold set at p < 0.05, unless otherwise stated.

Comparisons between groups were performed using a t-test (Figure 2a) or one-way ANOVA (Figures 3b, 5, and 8), or two-way ANOVA (Figures 6b and 7b), followed by Tukey’s post-hoc test with adjustment for multiple comparisons when applicable. Correlations between parameters (Figure 7b) were evaluated using Pearson’s correlation coefficient.

Data are presented as mean ± SD, except for Figure 7b, which is represented as mean ± SE. The distribution of morphological parameters in Figure 7b is provided in Supplementary Figure S6.

## Acknowledgements

The Authors acknowledge the contribution of Thomas Iskratsch and Emilie Mahuenda (SEMS-QMUL) for their advice on how to work with PDMS tuneable stiffnesses substrates, Alicia El Haj (Birmingham university) and Liisa Marketta Blowes (CREATElab QMUL) for their scientific advice and guidance and Erica Di Federico for her help with the PDMS mechanical characterisation.

Funders: This project was funded by the UK RMP - MRC (MR/T015462/1) and Dunhill Medical Charity (RPGF1802\23).

## Supplementary Figures

**Figure S1:**
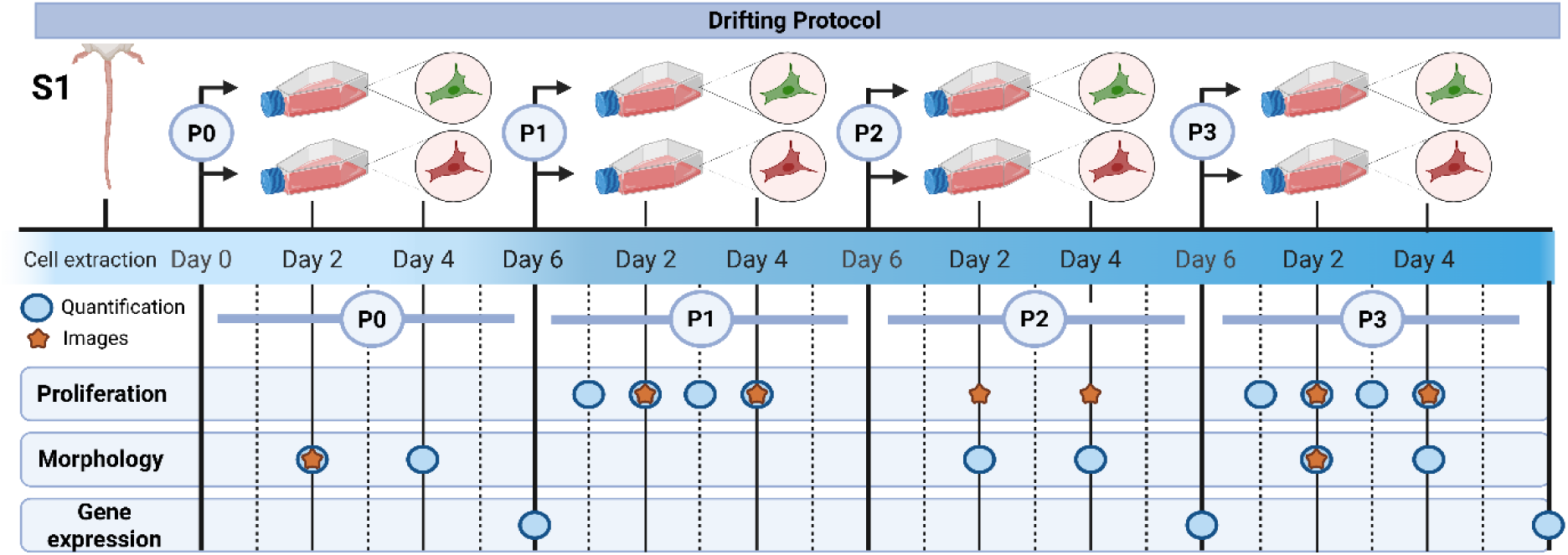
Timeline for cell phenotype drifting analysis. A schematic of the experimental protocol identifies the analysis time points selected for exploring IFM and FM cell phenotype drift though passage on TCP.

**Figure S2:**
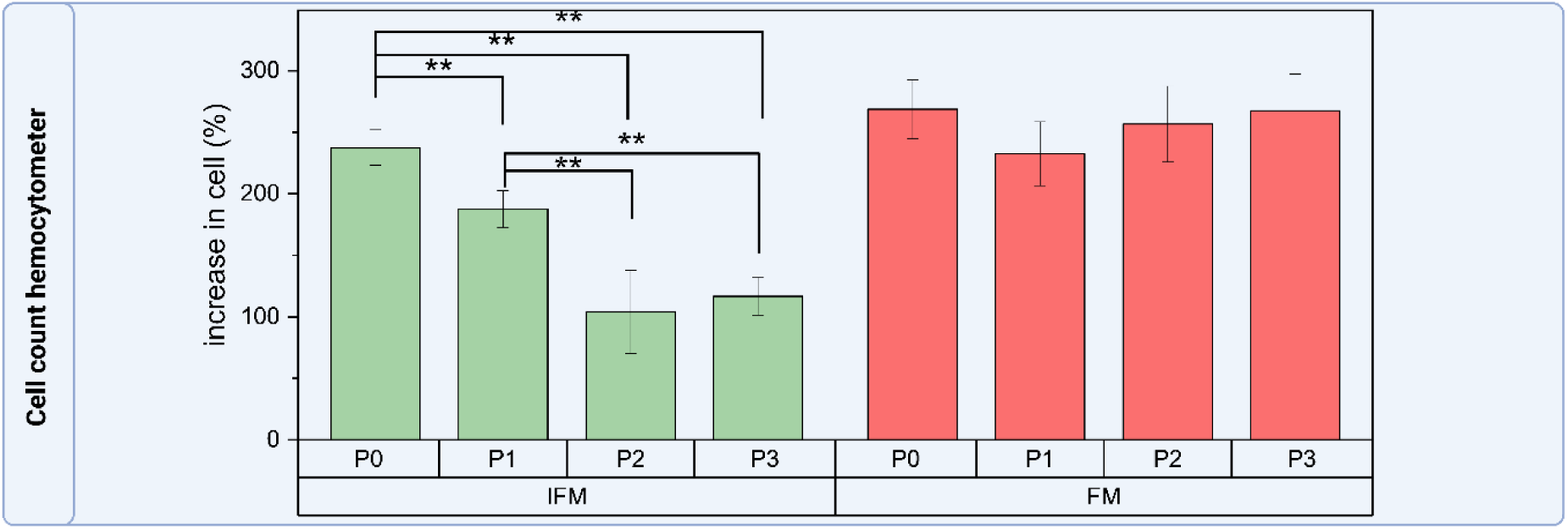
The increase in IFM cell numbers during each passage reduces with increasing passage number during culture on tissue culture plastic. Cell count data using haemocytometry shows the percentage increase in cell numbers from day 0 to 6 of culture at each passage. IFM and FM cells proliferated at a similar rate at P0 when first plated. However, following the first passage, IFM cell count was significantly reduced (p<0.01, n=4), and reduced even further in subsequent passages (p<0.001, n=4), showing a 93% reduction in proliferation rate from P1 to P3 (p=0.006).

**Figure S3:**
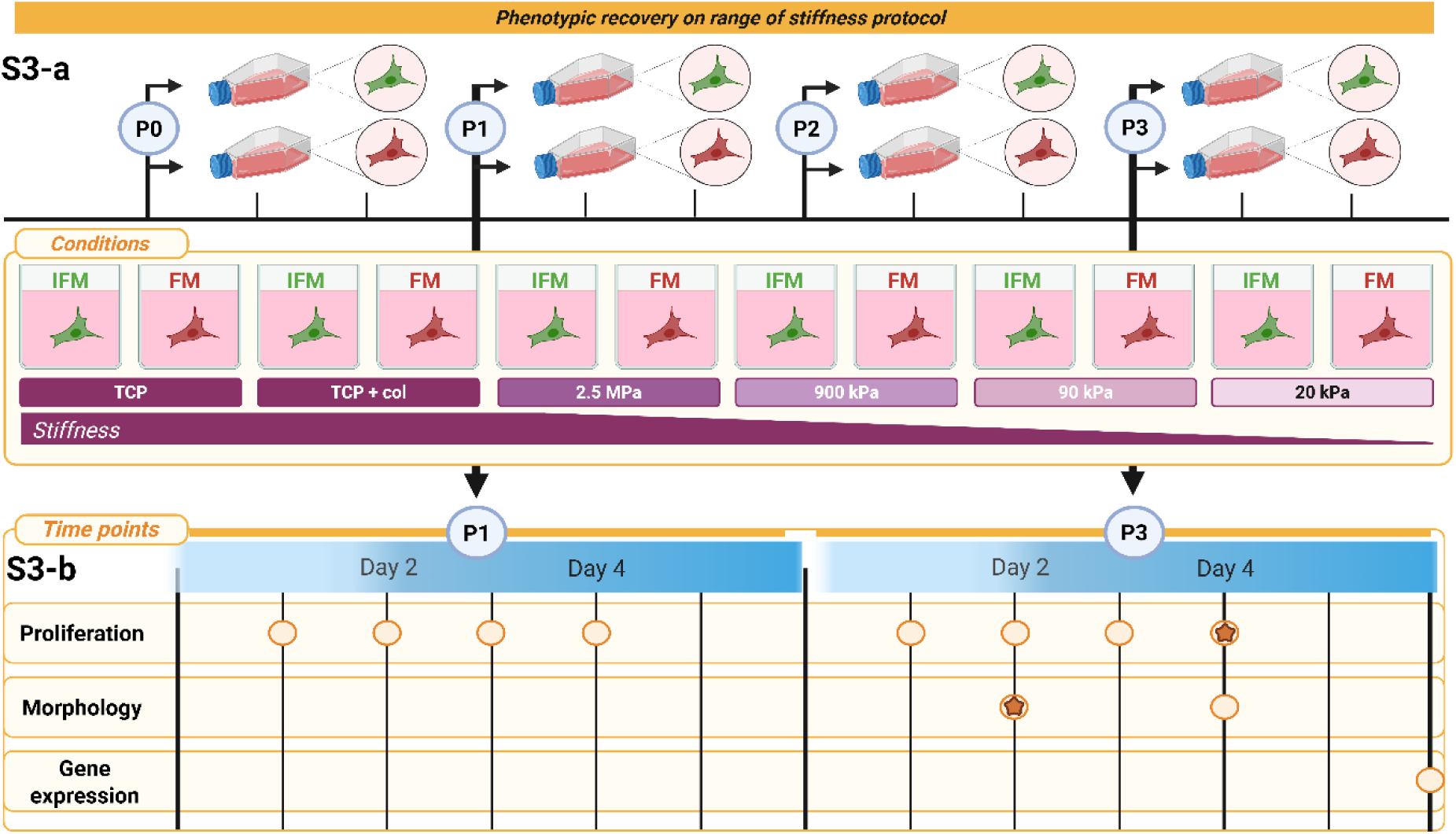
Timeline for cell phenotype recovery analysis. a) Schematic illustration of the protocol in which cells were cultured on TCP until either P1 or P3 at which point they were transferred to a range of softer substrates to explore the recovery of cell phenotype. b) The time points and passages selected for each analysis approach are outlined.

**Figure S4:**
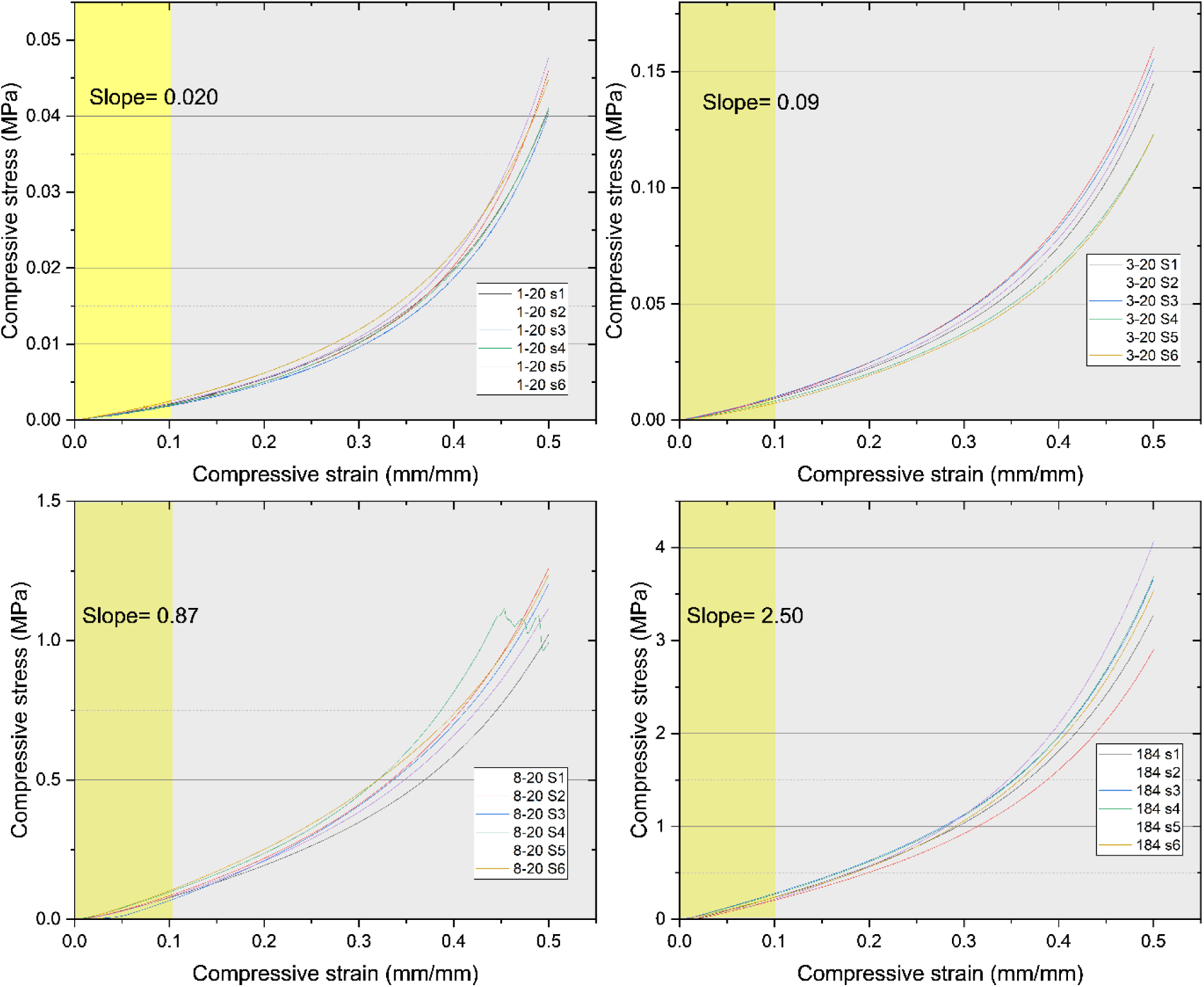
Compressive stress-strain behaviour of different PDMS concentrations. Compressive stress-strain curves for PDMS samples with varying compositions of PDMS 527 and PDMS 184. Samples tested include PDMS 527/PDMS 184 ratios of 1:20, 3:20, 8:20, and pure PDMS 184, with n=6 for each composition. The Young’s modulus for each sample was calculated from the gradient of the linear region of the stress-strain curve between 0–10% compression (shown in yellow).

**Figure S5:**
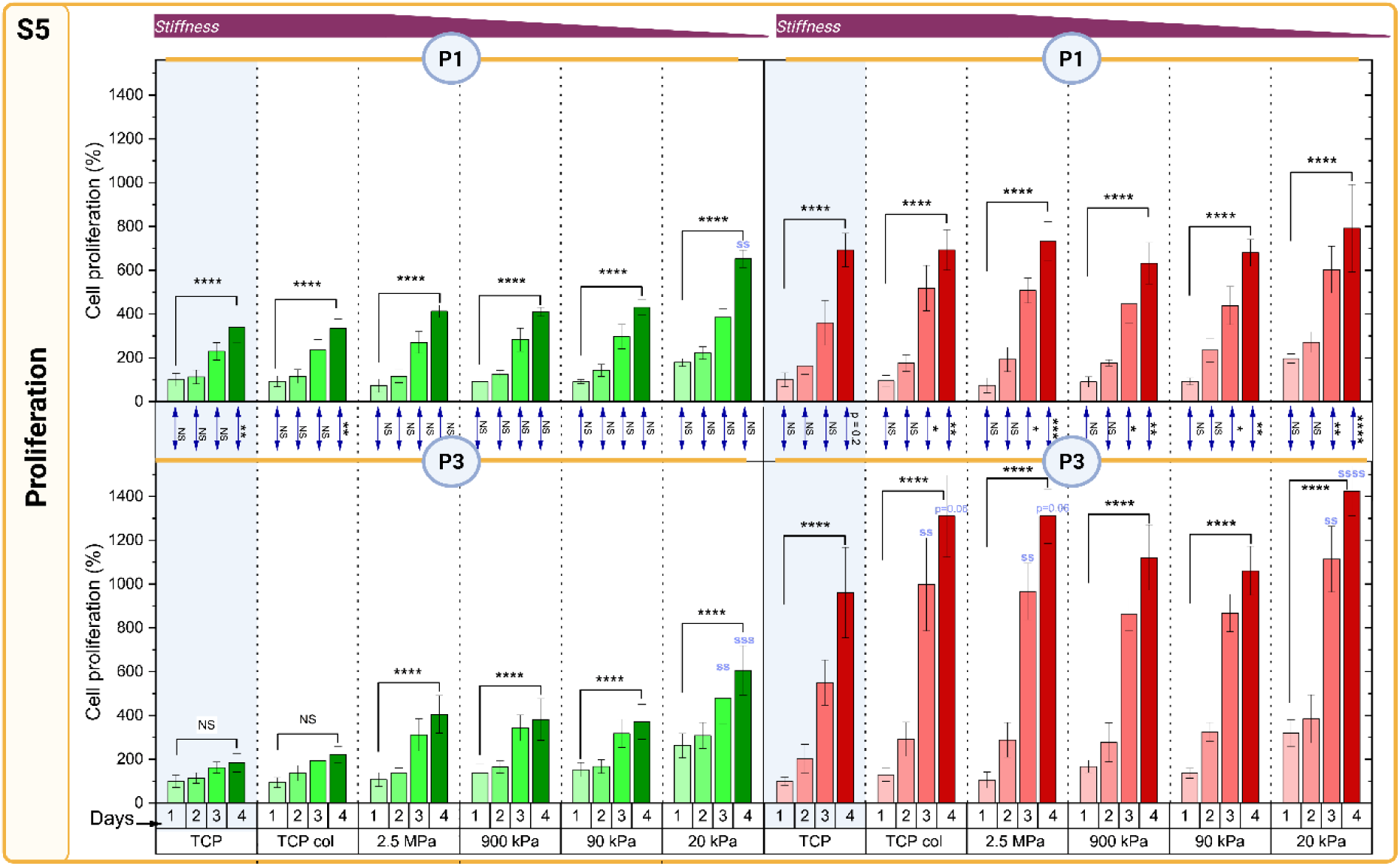
IFM cell proliferation recovers on softer surfaces. DNA quantification for cells cultured for 4 days across the range of stiffness highlights that IFM cell proliferation is higher on softer surfaces, and consistent whether cells are transferred at P1 (top row) or P3 (bottom row), whilst FM cell proliferation is unaffected by TCP culture or transfer to softer surfaces. DNA quantity is measured using Hoechst and normalised to day 1 (TCP) for each individual experiment. * highlights significant differences across days, $ highlights a significant difference in a stiffness condition relative to TCP. * or $ = p<0.05, ** or $$ = p< 0.01, *** or $$$ = p<0.001, **** = p<0.0001.

**Figure S6:**
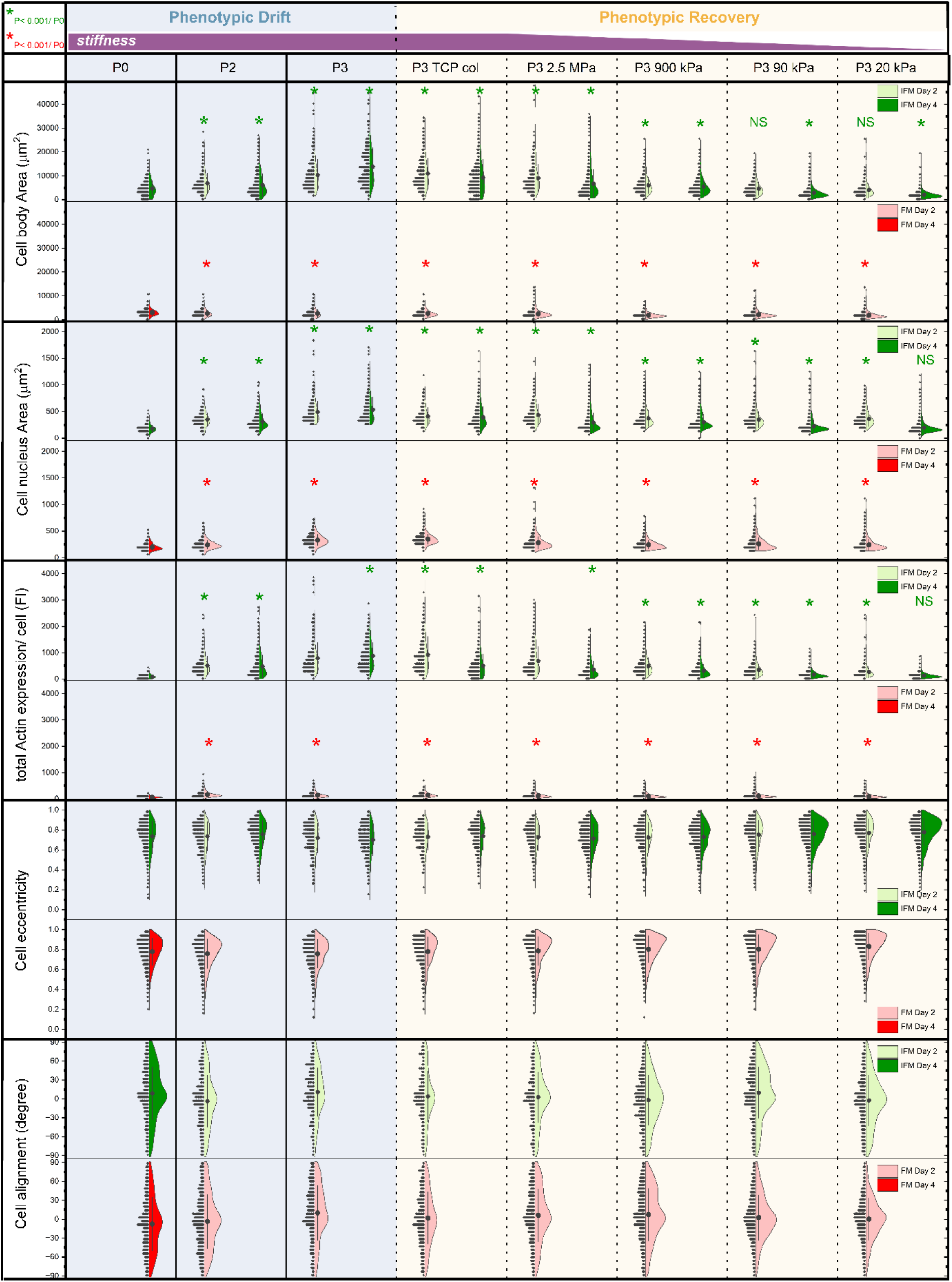
Morphological characteristics of IFM and FM cells in culture on TCP and PDMS substrates over passages. IFM-FM cell morphology parameters on varying substrate stiffnesses, showing cell body area, nucleus area, actin abundance/cell, cell eccentricity, and cell alignment, for IFM cells (green) and FM cells (red). Data is presented using the Kernel Density Estimation (KDE) method, with the right side of each violin plot representing the distribution and the left side showing the dataset repartition, for a minimum of 200 cells in each condition with error bars indicating standard error. Data on the left (blue background) depicts response from P0 to P2 when cells were cultured across passages on TCP. Data on the right (yellow background) depicts cell response when transferred onto a range of increasing softer substrates (1.5 MPa to 4–8 kPa) at the start of P3. Within each condition, day 2 of culture is shown in light green or light red and day 4 in dark green or dark red. Day 4 is missing in most FM cell conditions as cells were too confluent to accurately assess parameters. Day 2 at P0 is missing as cells were still settling. Statistical significance relative to the P0 TCP reference is indicated by * = p < 0.001. Data is summarised in figure 7a.

**Figure S7:**
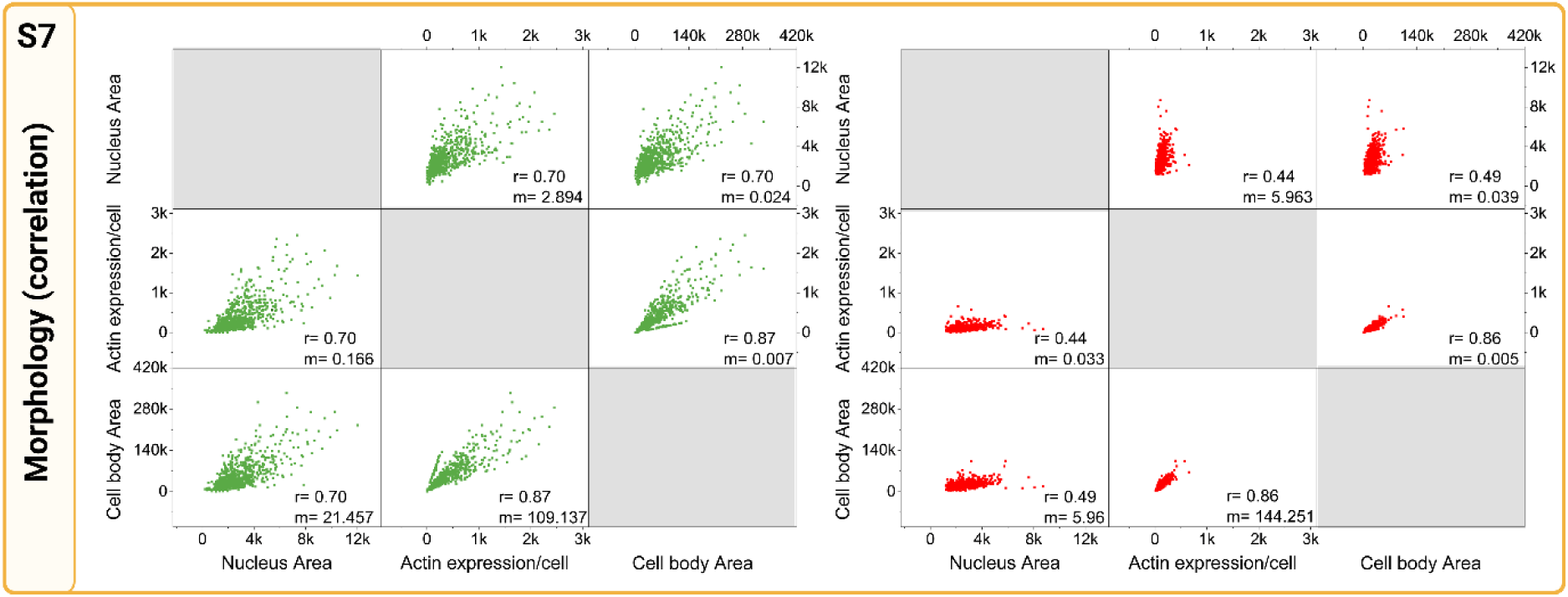
IFM cell morphology is strongly impacted by substrate stiffness. Correlations between cell body area, nucleus area and actin abundance/cell are shown for IFM cells (green) and FM cells (red) with the associated Pearson corelation coefficient (r) and the slope (m).

**Figure S8:**
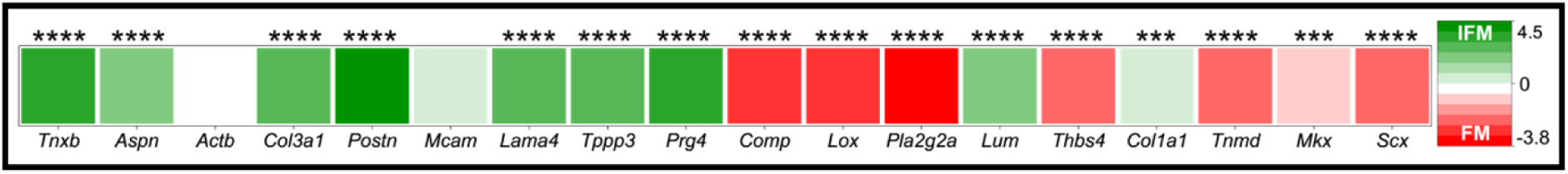
Heatmap of average gene expression values for the proposed IFM cell and FM cell clusters derived from equine SDFT single-cell RNA sequencing. (14). A calculation of the relative abundance of IFM and FM genes from equine single cell RNA sequencing data, calculated using the same method as adopted for the analysis of isolated rat IFM and FM cells in the current study (Figure 2) to facilitate comparison. Values represent the log₂ fold change of gene expression in IFM relative to FM, normalised to GAPDH. (*= p<0.05, **= p< 0.01, ***=p<0.001, ****= p<0.0001).

**Figure S9:**
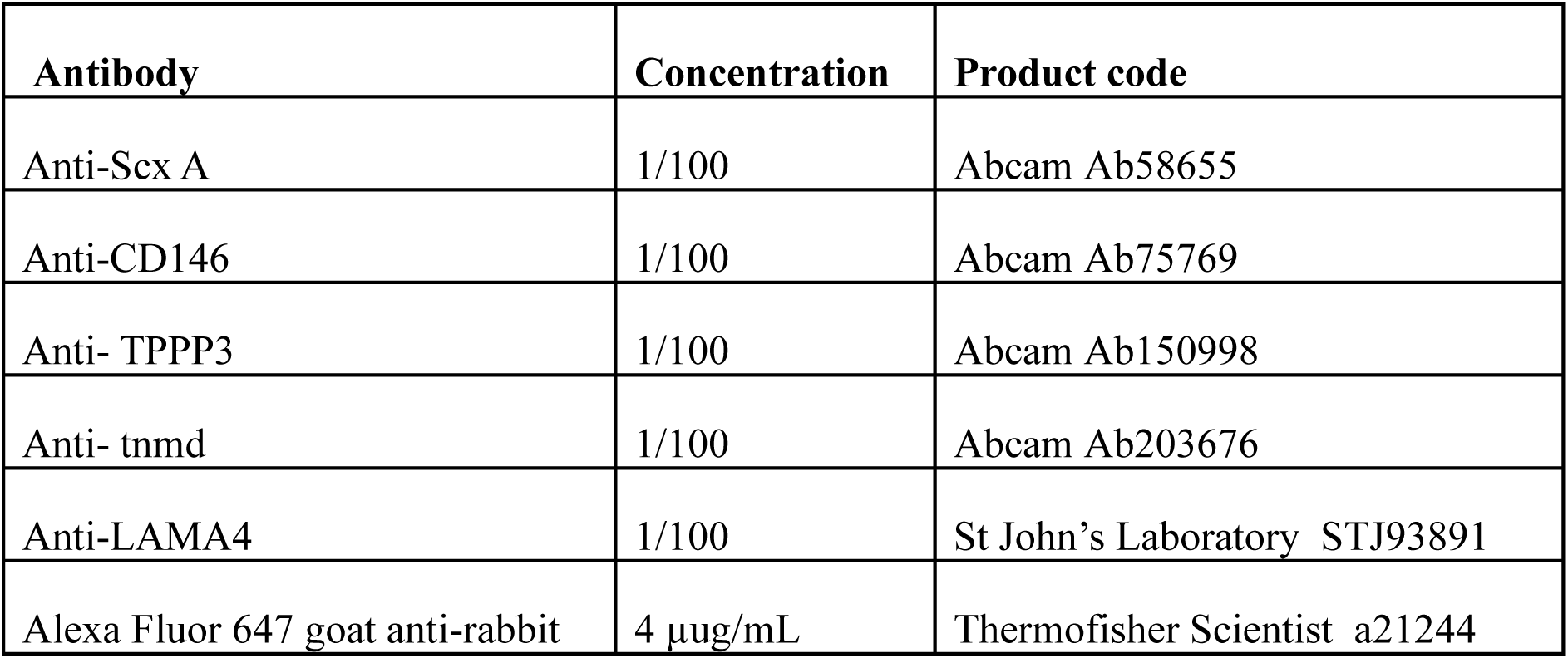
List of antibodies, concentrations and product code specifications used for immunofluorescence on rat tail tendons.

